# Expression based polygenic scores - A gene network perspective to capture individual differences in biological processes

**DOI:** 10.1101/2024.03.01.583008

**Authors:** Barbara Barth, Euclides José de Mendonça Filho, Danusa Mar Arcego, Irina Pokhvisneva, Michael J. Meaney, Patrícia Pelufo Silveira

## Abstract

Incorporating functional aspects into polygenic scores may accelerate early diagnosis and the discovery of therapeutic targets. Yet, existing polygenic scores summarize information from genome wide statistical associations between SNPs and phenotypes. We developed the novel biologically informed, expression-based polygenic scores (ePRS or ePGS). The method characterizes tissue specific gene co-expression networks from genome-wide RNA sequencing data and incorporates this information into polygenic scores. Performance and characteristics of the ePGS were compared to traditional polygenic risk score (PRS). We observed that ePGS differs from PRS for aggregating information on; i. the relation between different genes (co-expression); ii. the levels of tissue-specific gene expression; iii. the genetic variation of the target sample; iv. the tissue-specific effect size of the association between genotyping and gene expression; v. the portability across different ancestries. Variations in the ePGS represent individual variations in the expression of a tissue-specific gene co-expression network, and this methodology may profoundly influence the way we study human disease biology.

## Main

Genome wide association studies (GWAS) are used to identify genetic variants statistically associated with a disease or trait^1^ by comparing single nucleotide polymorphisms (SNPs) across the genome in cases and controls. An initial objective was to identify individual common variants closely linked to phenotype that might account for a substantial portion of inter-individual variation. However, it is now clear that common disorders and complex traits are instead highly polygenic, reflecting the influence of thousands of polymorphisms, each with relatively small effects. Polygenicity led to the development of polygenic risk scores (PRSs) that are calculated from GWAS results in target samples to reflect a cumulative influence of risk alleles. PRS aggregates the GWAS information by summing the risk alleles count weighted by the effect size for each SNP presented in the GWAS ^2,3^. PRS combines the isolated small effects of multiple genetic variants in a single score that represents the genetic risk for a disease or variation in the expression of a trait. The use of PRSs has proven effective in defining main effects of heritable genetic variations in relation to a wide range of outcomes. Moreover, PRSs are a continuous measure that offer a plausible alternative to candidate gene approaches.

Polygenicity involves the function of diverse genes and molecules that interact with each other in cellular networks^4^. Genes do not operate in isolation but conjointly in tissue-specific networks that regulate molecular events and precise biological functions^5^. A gene network involves a number of genes co-expressed within a specific tissue or brain region that exert a concerted effect on a target biological process. Since they rely solely on DNA sequence variation, existing PRS methods do not capture these biological intricacies and functional relations of tissue-specific gene networks. The challenge was to create a genomic metric that would reflect the influence of genetic variation, as does the PRS method, but do so within the context of a tissue-specific gene network. To meet this challenge, we created an innovative approach to genomic profiling that characterizes gene networks based on the levels of co-expression within a specific tissue^6–17^. The co-expression based polygenic score (ePRS or ePGS) method integrates information from both GWAS and tissue-specific RNA sequencing (RNAseq) data sets.

In the examples presented here using the ePGS technique, we focus on specific brain regions, but the method can be applied to any tissue. There are two approaches to the definition of the co-expression networks that depend upon the research objective. One approach is designed to test specific hypothesis regarding the function of a specific gene network in a specific brain region. In this instance (see **Figure 1**) a gene network is constructed by focusing on a target gene in a specific brain region. In a series of studies, we focused on dopamine signaling in the prefrontal cortex and thus created a co-expression network comprised of genes in which the expression is significantly (i.e., r>0.5) correlated with that of *SLC6A3*, which encodes the dopamine transporter. For the sake of comparison, we created *SLC6A3*-based co-expression networks from RNAseq data sets in an alternative brain region. This approach allows the researcher to define the region-specificity for any outcomes. A virtue of this approach is the ability to test hypotheses, often derived from studies with model systems, using human data sets. The second approach is aligned to discovery and employs Whole-Genome Co-Expression Network Analysis (WGCNA)^18^ to identify co-expression modules from RNAseq data. The resulting modules can then be statistically tested for associations with treatment or traits of interest. The module statistically related to the trait of interest then serves as the gene network for the calculation of the ePGS.

**Figure 1.**
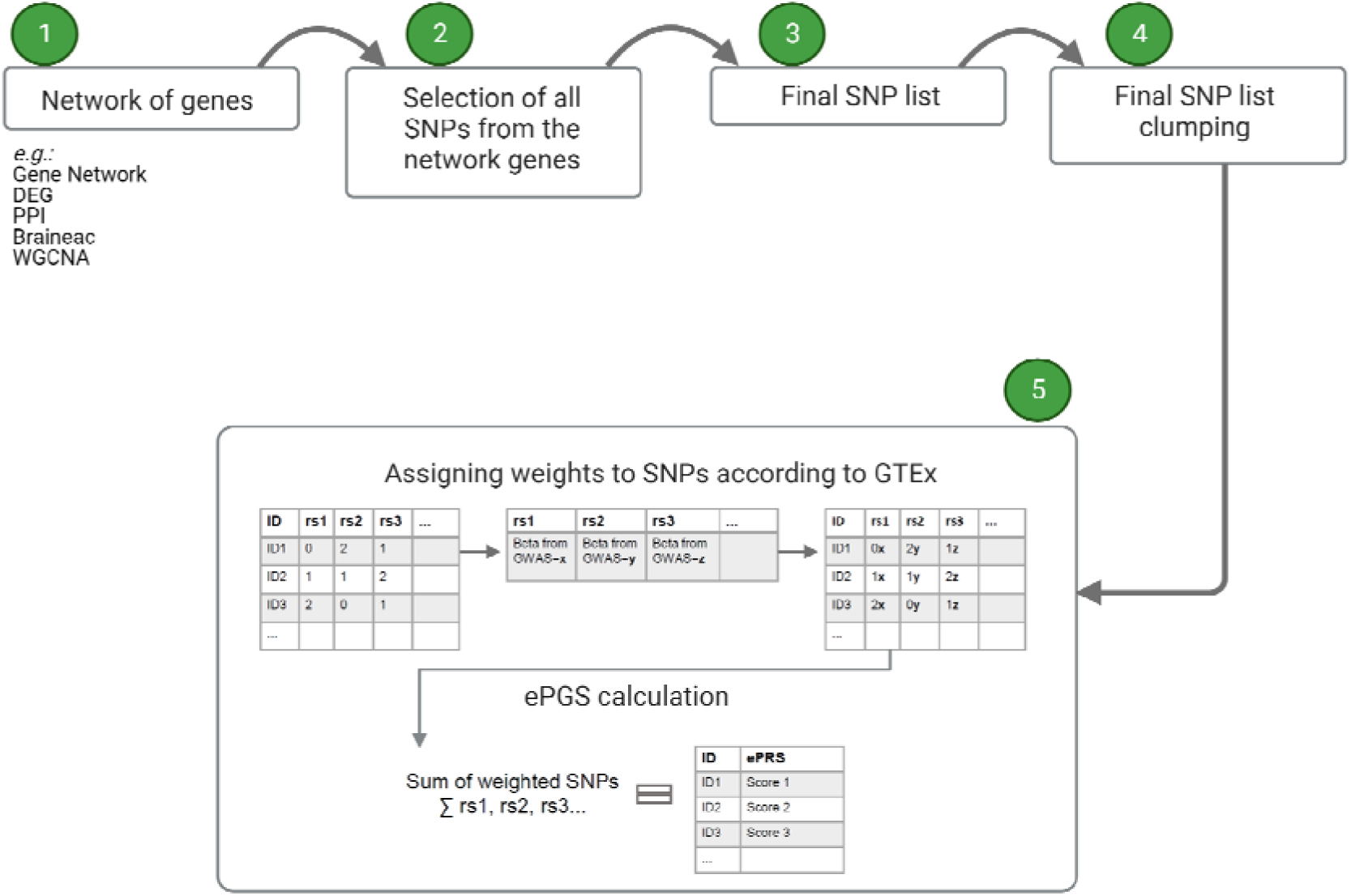
Schematic figure representing the key steps to calculate the ePGS. **1)** Construction of a network of genes that is defined by a set of genes that interact in a biologically meaningful way. Some examples are co-expression of transcripts from animal models (GeneNetwork), as used in the current study, and different expression analysis (DEG). Additionally, it can be defined by protein-protein interaction (PPI), co-expression of transcripts from human samples (Braineac) and by weighted gene co-expression network analysis (WGCNA). At this step, tissue specificity can be defined by selecting transcript data from specific tissues of interest. The list of genes can also be filtered by a specific developmental time point, for example, by using publicly available databases such as the BrainSpan^34^. Furthermore, the list of genes can be filtered by other conditions and interests. **2)** Selection of all existing SNPs from the gene network was done using biomaRt package. From this list we retained common SNPs with a) SNPs from the study sample genotyping data and b) SNPs present in GTEx (which is a genome-wide analysis that has gene expression as the outcome; GTEx was chosen to weight the selected SNPs in the examples provided here). The common SNPs represent the final SNP list that is subjected to linkage disequilibrium clumping (r^2^>0.2). **5)** Weight the SNPs: the number of effect alleles (genotype information from the study sample) at a given SNP is multiplied by the effect size of the association between SNPs and the gene expression (GTEx). The sum of all weighted SNPs for each individual corresponds to the individual ePGS.

The gene network of interest then serves as the basis for the selection of genes used in the formulation of the ePGS. SNPs from these genes are functionally annotated and subjected to linkage disequilibrium clumping for removal of highly correlated SNPs. A count function of the number of effect alleles at a given SNP is established and weighted by the effect size of the association between the individual SNP and the expression of the related gene in a specified tissue using the Gene Tissue Expression (GTEx ^19^) human RNAseq data sets. The sum of these values from the total number of SNPs defines the ePGS at the level of the individual subject (**Figure 1, Supplemental Figure 2**) (**Supplemental Methods**).

The ePGS combines information on: i. the relation between different genes (co-expression); ii. the levels of tissue-specific gene expression (bulk or single-cell genome wide RNAseq); iii. the genetic variation of the target sample (genotyping data); iv. the tissue-specific effect size of the association between variants and gene expression (GTEx). Therefore, variations in the ePGS represent individual variations in the genetically-determined capacity for the expression of the genes that comprise the tissue-specific gene co-expression network. In this paper we present the ePGS technique, its method of calculation and compare its features and score content with a traditional PRS.

## Results

### Expression-based polygenic scores (ePGS) calculation

The steps by which an ePGS is constructed are summarized in **Figure 1**. We first describe the methods for the identification of tissue-specific gene networks, which are the essential feature of the ePGS approach. Researchers can use both co-expression^6–16^ and differential expression^17^ data, from publicly available or their own datasets, see **Supplementary Figure 2**. Publicly available data sets include RNAseq databases for both rodents (e.g. GeneNetwork ^20^) and humans (e.g. BrainEAC ^21^) that can be used to identify gene networks. In the examples presented here we focus on a specific brain region, but the method can be applied to any tissue.

Since expression of gene networks vary from region to region, obtaining gene networks that are tissue specific informs on the relevance of both the gene network and the brain region or tissue. A formidable advantage of the ePGS approach is the ability to create a genomic metric by which to test hypothesis concerning tissue-specific gene expression profiles in any human data sets for which there is both genotyping and the target phenotypic measure. In this instance a gene network is constructed by focusing on a target gene in a specific brain region. For the examples that will be discussed here, we have focused on dopamine signaling in the mesocortical pathway, more specifically, the prefrontal cortex (PFC), the final target of this pathway. To achieve this, we constructed a co-expression network comprising genes whose expression is notably correlated (i.e., r ≥ 0.5) in the PFC with either SLC6A3, responsible for encoding the dopamine transporter, or with the dopamine receptor D2 gene (DRD2), two important regulators of dopamine neurotransmission in the brain (See **Supplemental Table 1**) (see **Figure 1** for schematic representation and **Supplemental Figure 2** for gene co-expression rationale). The calculations were performed separately for each gene network of interest using the GeneNetwork (http://genenetwork.org) database from RNAseq data from mice. Note, the cut-off for the correlation coefficient is arbitrary, based on conventionally regarded as moderate to high correlation. For the sake of comparison and to establish tissue specificity, we create a co-expression network from RNAseq data sets in an alternative brain region. This feature allows the researcher to statistically establish associations that are tissue or brain region specific.

When the identification of the gene network is anchored to a specific target gene, the gene network is composed of the genes significantly co-expressed with that target gene in a specific brain region or tissue (**Figure 1**). Using biomaRT R package^22,23^ (Ensembl GRCh37) the co-expressed genes are converted to human homologous genes, and all the existing SNPs from these genes are gathered. Common SNPs were selected between the three sources (the SNPs gathered from the gene networks of interest, the SNPs from the GTEx project^19^ data in human PFC and with the SNPs from the study sample (1000 Genomes Project^24^)) and were subjected to linkage disequilibrium clumping (r^2^<0.2) within 500kb radius, to inform the removal of highly correlated SNPs. The number of effect alleles at a given SNP is weighted using the estimated effect of the tissue specific genotype-gene expression association from the GTEx project^19^. We also accounted for the direction of the co-expression of each gene with *SLC6A3* or *DRD2* by multiplying the weight by -1 in case the expression of a gene was negatively correlated with the expression of the *SLC6A3* or *DRD2* genes. The sum of the weighted values from all SNPs, divided by the number of SNPs, provided the region-specific ePGS scores.

The ePGS scores were calculated separately for each ancestry in the 1000 Genomes Project, which includes African (N=661), American (N=347), East Asian (N=504), European (N=503) and South Asian (N=489). Since the majority of donors in the GTEx project were of European ancestry^25^ (see donor information at: https://gtexportal.org/home/tissueSummaryPage), most of the comparisons demonstrated here used 1000 Genomes Project European sample, for both ePGS and PRS (see Supplemental material, the exception being the analysis comparing the scores across all ancestries). The *SLC6A3* network for European ancestry included 262 genes and 15387 SNPs. The *DRD2* network for European ancestry had 281 genes and 12595 SNPs (See **Supplemental Table 1** for a description of genes and SNPs included in all scores described in the study).

### ePGSs reflect cohesive, biologically meaningful gene networks

We then compared the gene network structure represented by same size ePGS and PRS. To achieve that, we mined gene co-expression information from GeneMANIA^26,27^ (http://genemania.org) to identify and quantify connections between the genes from each score. GeneMANIA provides coexpression information between all genes from a queried gene list. We also used the Centiscape tool^28^ in Cytoscape®^29^, to estimate two centrality measures of the networks: degree, which is the number of connections between each node (each gene) and betweenness, that estimates the number of times a node lies on the shortest path between other nodes. **Figure 2a** depicts the gene network for *SLC6A3* PFC ePGS (number of genes = 262), with a dense connection pattern between genes. Similar sized PRSs for broad depression resulted in a network, depicted in **Figure 2b** (number of genes = 265). When comparing the total degree between genes in the different scores using a one-way ANOVA, results show that the *SLC6A3* PFC ePGS derived gene network has significantly more total connections than the broad depression PRS (**Figure 2c**). The same results were found for the *DRD2* PFC ePGS (281 genes, **Supplemental Figure 1a**) and its comparable size broad depression PRS (**Supplemental Figures 1b and 1c**).

**Figure 2.**
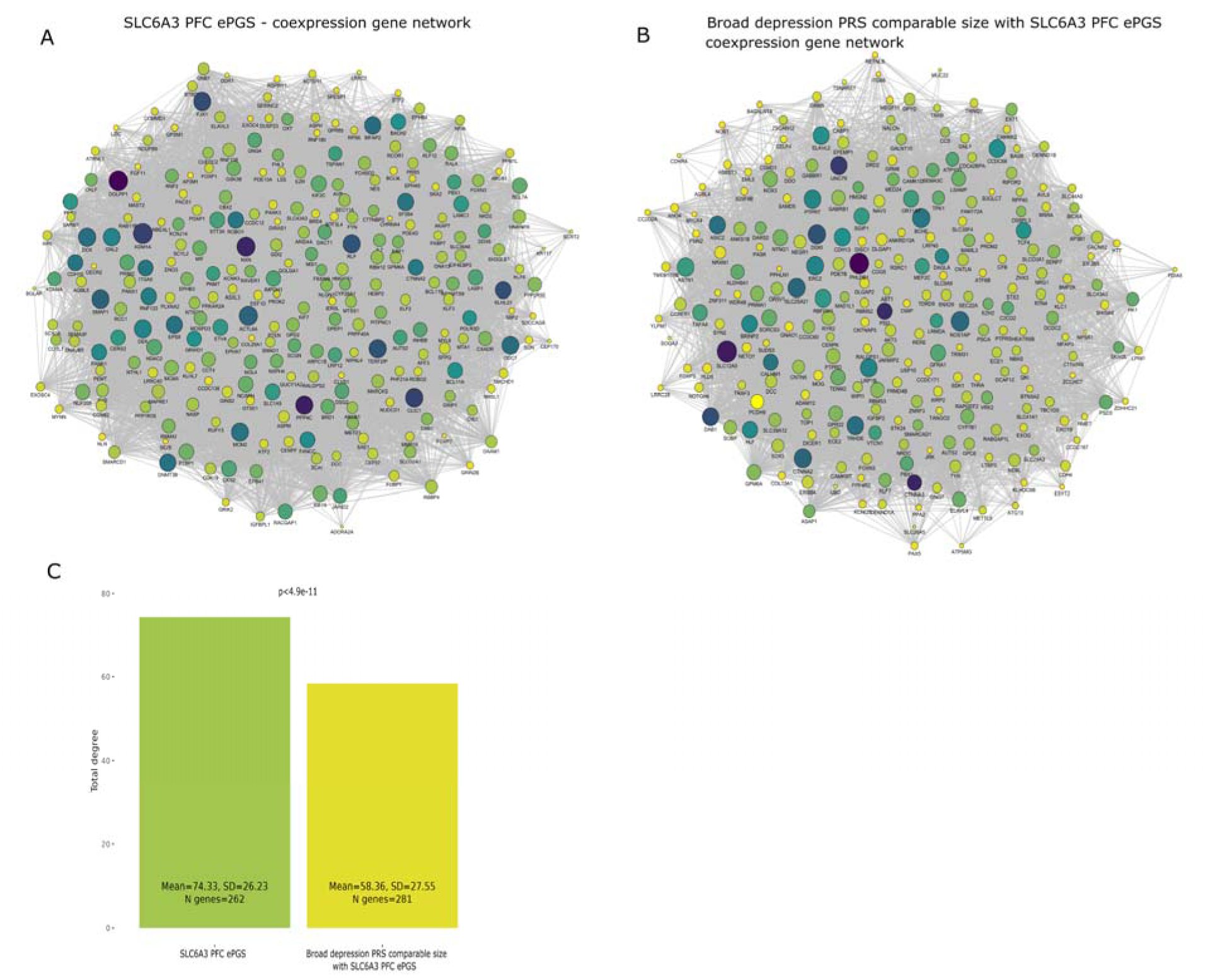
Network visualization comparison of *SLC6A3* derived ePGS and comparable size PRSes. **a)** *SLC6A3* PFC ePGS gene network; **b)** Broad depression PRS gene network comparable size with *SLC6A3* PFC ePGS; **c)** One-way ANOVA of total connectivity (total degree values) for ePGS and PRS comparable size. Gene co-expression interactions were obtained from GeneMANIA (http://genemania.org) and used to generate the networks with Cytoscape® application, which specifies amount of interactions between pairs of genes based on their co-expression, represented by the number of edges (gray lines) in the networks. The Centiscape plug-in in Cytoscape® was used to calculate the centrality of the genes in each network, defining the degree (number of connections with other nodes, represented by node size, in which bigger nodes indicates more connections with other nodes) and betweenness (number o times a node lies on the shortest path between other nodes, represented by node’s color in which darker colors indicate higher betweenness in the networks) for the components of the networks.

It is important to highlight main conceptual differences between ePGS and PRS that can explain dissimilarities in total connectivity. PRSs are built selecting SNPs from a GWAS based on their genome-wide significance level, and for that reason both intron and exon DNA sequences are considered. Introns are non-coding DNA sequences within the genome, and therefore are not mapped to genes. Introns embody 25% of the human genome and are 4 to 5 times the size of exons^30^. In fact, a large number of significant SNPs from GWAS are in intronic and intergenic regions^31,32^. On the other hand, the ePGS is built from gene co-expression information, and therefore considers only protein-coding DNA sequences, the exons, resulting in every SNP being mapped to a gene. The ePGS maps into a dense group of genes (higher connectivity) that interact with each other, possibly representing associated molecular functions as described below.

### ePGS and PRS represent different biological mechanisms

Because of the differences in SNP selection between ePGS (a gene co-expression network identified in RNAseq data) and PRS (statistically significant SNPs from a GWAS), it is expected that the two scores will differ in the biological mechanisms that they represent. We compared PRS and ePGS enrichment analyses using MetaCore™ (Clarivate Analytics, version 21.4) (https://portal.genego.com) and the function “compare experiments”. We identified a significant common gene ontology (GO) term and exported unique elements from each network that are significantly associated to that GO term (FDR < 0.05) for comparison purposes. Networks were constructed for direct interactions between selected objects and filtered for brain tissue and human species.

It is noteworthy that “neuron differentiation (FDR<0.001)” was a common GO process associated with genes from both PRSs and ePGS genes. However, this finding was due to different element networks in each score (**Figure 3**). In ePGS, “neuron differentiation” was mapped to elements such as “Nestin”, which is present in neural stem and progenitor cells and directly involved in differentiation process^33^. In PRS, “neuron differentiation” was mapped to elements such as “olfactory receptor” and less connections are seen between elements. Taken together, the findings depicted in **Figure 3** suggest that while both ePGS and PRSs are linked to processes related to neuron projection development, these relations occur via unique and specific mechanisms. The unique elements related to the ePGS score, in these examples, are richer and more connected, suggesting that variations in the ePGS score possibly represent variation on these specific biological processes.

**Figure 3.**
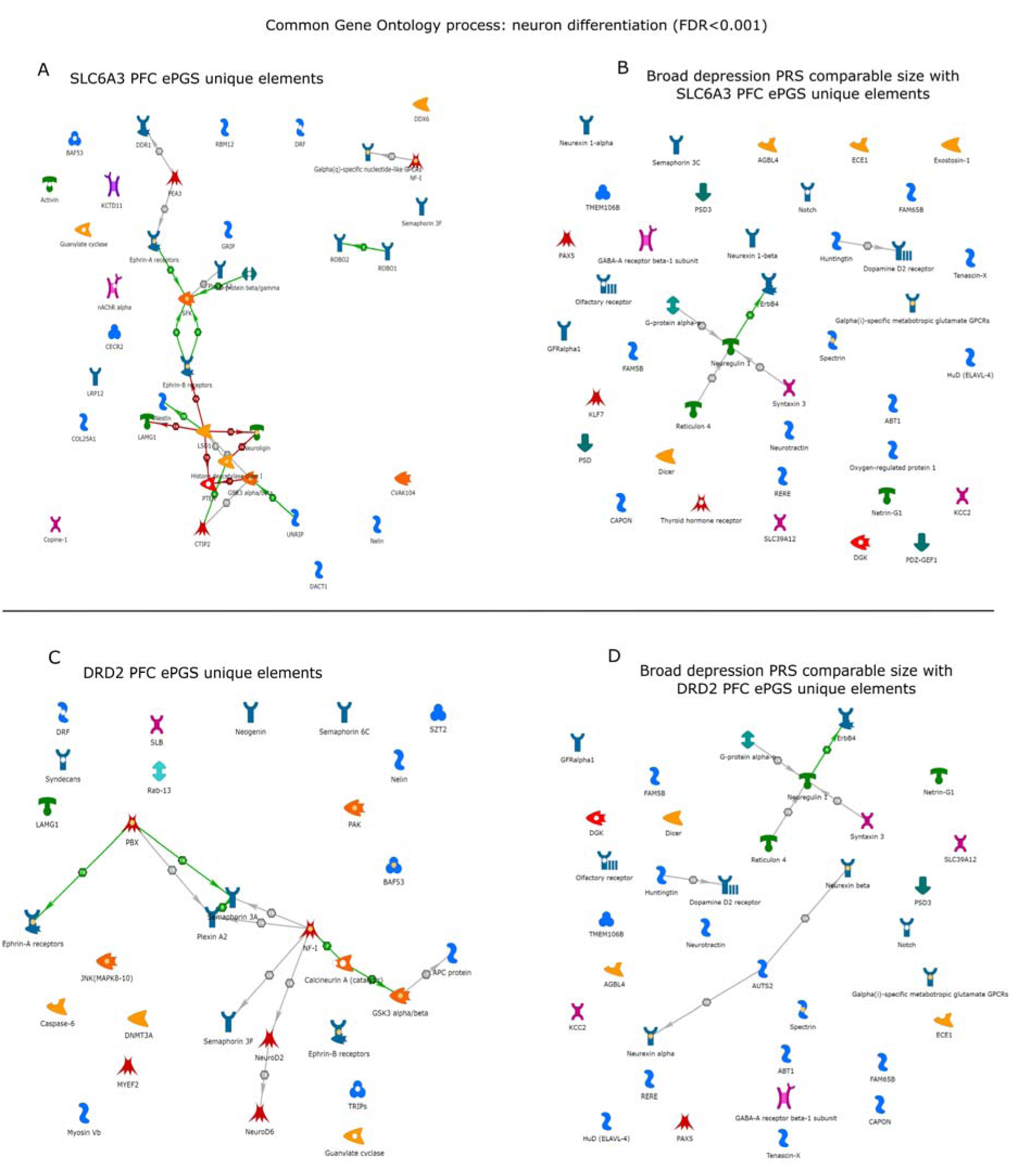
Unique elements for ‘neuron differentiation’, a common gene ontology enrichment analysis term for both ePGS and PRS. Gene ontology (GO) enrichment analysis was performed using Metacore®. The function “compare experiments” was used to obtain common significant (FDR <0.05) GO terms between the gene networks while also identifying the unique elements from each network that are significantly associated to the GO term. Networks were plotted in MetaCore® using the unique elements of each network for the GO enrichment term selected. Figures **a, b, c,** and **d** show visual comparisons of the different contributions of ePGS and PRS to the GO term. The details of the legends of the network’s figures can be found in https://portal.genego.com/legends/MetaCoreQuickReferenceGuide.pdf.

### ePGS genes represent co-expression networks that are preserved across species

Since our example ePGSs were originally informed by co-expression networks identified in mice (**Supplemental Methods**), we examined whether ePGS genes would also represent co-expression networks in humans and compare brain co-expression patterns between ePGS genes and traditional PRS genes. We used PFC gene expression data in human post-mortem brain tissue from the BrainSpan database (from embryonic to adulthood, N= 42)^34^ and analyzed the correlation between the expression levels in the PFC for the ePGS and PRS gene lists. It is important to note that in this comparison the gene list used for the ePGS originates from mouse, whereas that for the PRS is from human data sets. Our results show that ePGS gene networks, in the examples given here, have greater PFC gene co-expression percentage in humans in comparison to PRS gene lists (**Figure 4**). For the *SLC6A3* PFC ePGS, 40% of the gene pairs had an absolute expression correlation r>=0.5 and 80% of the correlations were significant at P<0.05. However, when using the genes of a traditional PRS for broad depression, a lower percentage of co-expression was observed with 17% of the gene pairs had an absolute expression correlation r>= 0.5 and only 62% of the correlations were significant at P<0.05. The same comparisons were done for the *DRD2* PFC ePGS and its respective comparable size broad depression PRS, and more robust co-expression patterns were consistently observed in ePGS in comparison to PRSs for broad depression (see **Figure 4**). The results from these examples indicate that ePGSs informed by mice RNAseq data represent brain gene co-expression networks also in humans, and these gene networks are more tightly connected than those represented by genes that constitute the traditional PRS in the examples seen here. This finding demonstrates a successful cross species translation of genome functional annotation into the ePGS scores.

**Figure 4.**
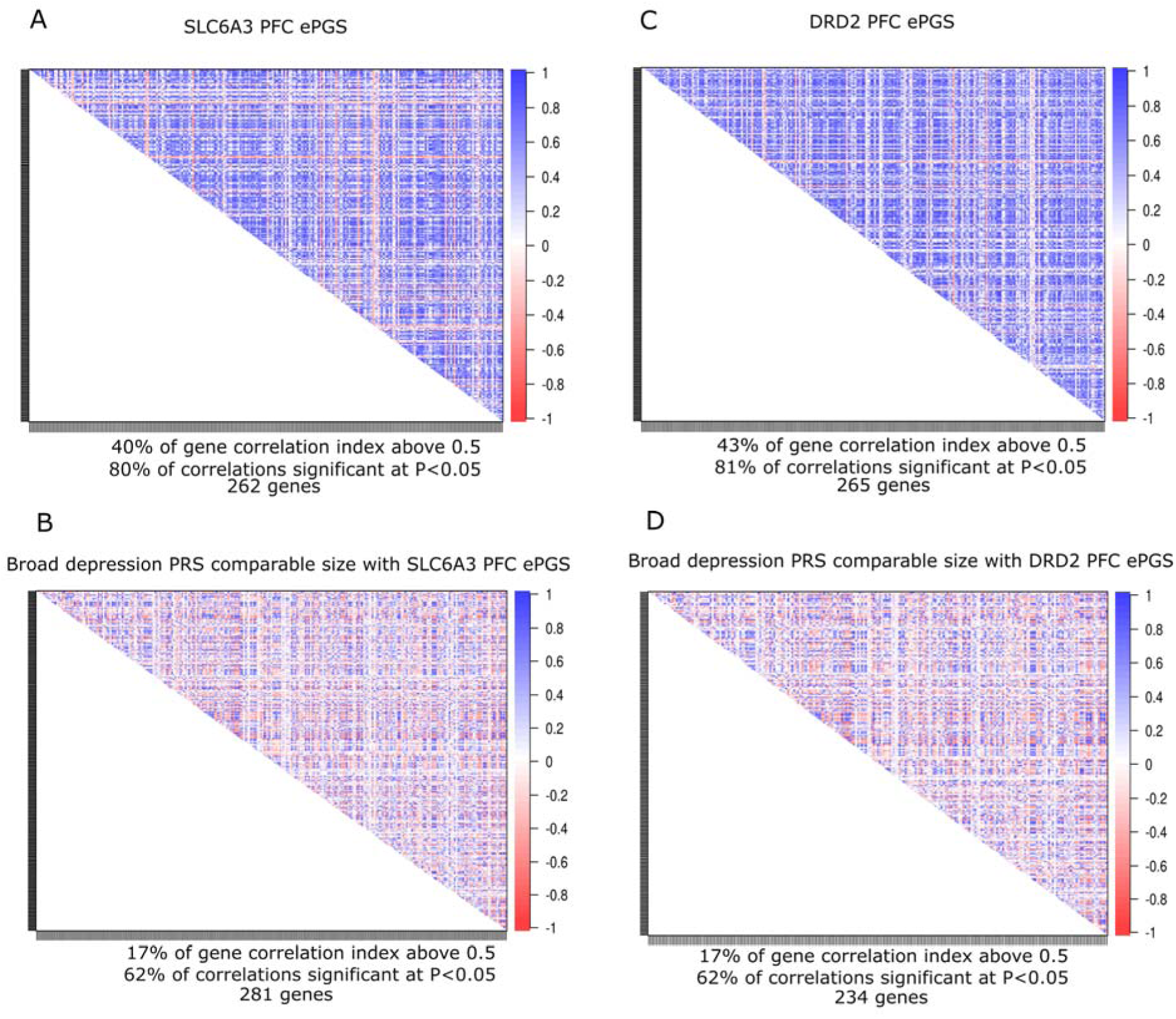
Correlation matrix of gene expression for ePGS gene networks and PRS gene networks based on BrainSpan human post-mortem brain tissue (from embryonic to adulthood, N=42). **a)** *SLC6A3* PFC ePGS gene network: 40% of the gene correlations was above 0.5 and 80% of the correlations are significant at P<0.05; **b)** Broad depression PRS gene network comparable size with the *SLC6A3* PFC ePGS: 17% of the gene correlations above 0.5; 62% correlations significant at P<0.05; **c)** *DRD2* PFC ePGS gene network: 43% of gene correlations above 0.5 and 81% of correlations significant at P<0.05; **d)** Broad depression PRS gene network comparable size with the *DRD2* PFC ePGS: 17% of the gene correlations above 0.5; 62% correlations significant at P<0.05.

### ePGS reflects tissue specific co-expression networks

The ePGS calculation is informed by RNAseq data, which quantifies genome-wide tissue-specific transcription (**Supplemental figure 2**). Therefore, the ePGS is based on tissue-specific gene co-expression data to identify the gene network. The tissue-specific genotype-gene expression association from GTEx is then to weight the ePGS SNPs. Thus, both the selection of the genes and their weighting are derived from tissue specific data sets. In contrast, a PRS is based on the genotype, which is the same across different cells and tissue types.

To exemplify the importance of tissue specificity, we compared two gene networks built on the same gene as the initial anchor, *SLC6A3*, in the PFC and the striatum. Please note the differences in visualization of the *SLC6A3* PFC (total number of genes = 262) and *SLC6A3* Striatum (total number of genes = 346) networks (**Supplemental Figure 3a**). We identified 53 genes in common between the networks (**Supplemental figure 3b**), which represents a small percentage of the total number of genes from both regions (21% for *SLC6A3* PFC ePGS and 15% for *SLC6A3* Striatum ePGS). This finding highlights the considerable tissue specificity of the networks, even when based on the same initial gene as the anchor, which demonstrates the ability of the ePGS to represent tissue specific information^35^.

### ePGS interacts with environmental variation

Despite a broadly-held conviction that genotype – phenotype relations can be context specific, the demonstration of gene x environment interactions has been controversial. The controversy was focused largely on candidate gene approaches that commonly failed to replicate and generally flew in the face of the polygenic nature of the target phenotypes. Unfortunately, despite its polygenic nature, investigations using polygenic scores derived from GWASs show only modest success in revealing gene-environment interactions^36^ ^37^ ^38^. This is actually unsurprising. A PRS is based on a GWAS using the most significant SNPs representing genetic variants strongly associated with a condition or trait. The considerable strength of the PRS method is the ability to capture polygenetically-determined predispositions for phenotypic outcomes as simple main effects using a continuous measure. A PRS is thus an ideal tool for the study of main effects of genetic variation. However, the reliance on SNPs that pass a designated level of statistical association with the phenotype of interest biases in favor of those variants that exhibit minimal environmental dependency. The implication is that SNPs in genes that that are highly dependent upon environmental context are less likely to emerge as significant as main effects in a GWAS, considering the rigorous GWAS-level of statistical significance for main effects. **Figure 5** shows a Manhattan plot for the broad depression GWAS^39^. SNPs in green are those included in the *SLC6A3* PFC ePGS, demonstrating that the variants included in the ePGS lie well below the GWAS significance level. This difference would be expected if SNP’s comprising an ePGS are context dependent. This may explain why the ePGS may be more suited to identify GxE interaction effects^40^ as documented below.

**Figure 5.**
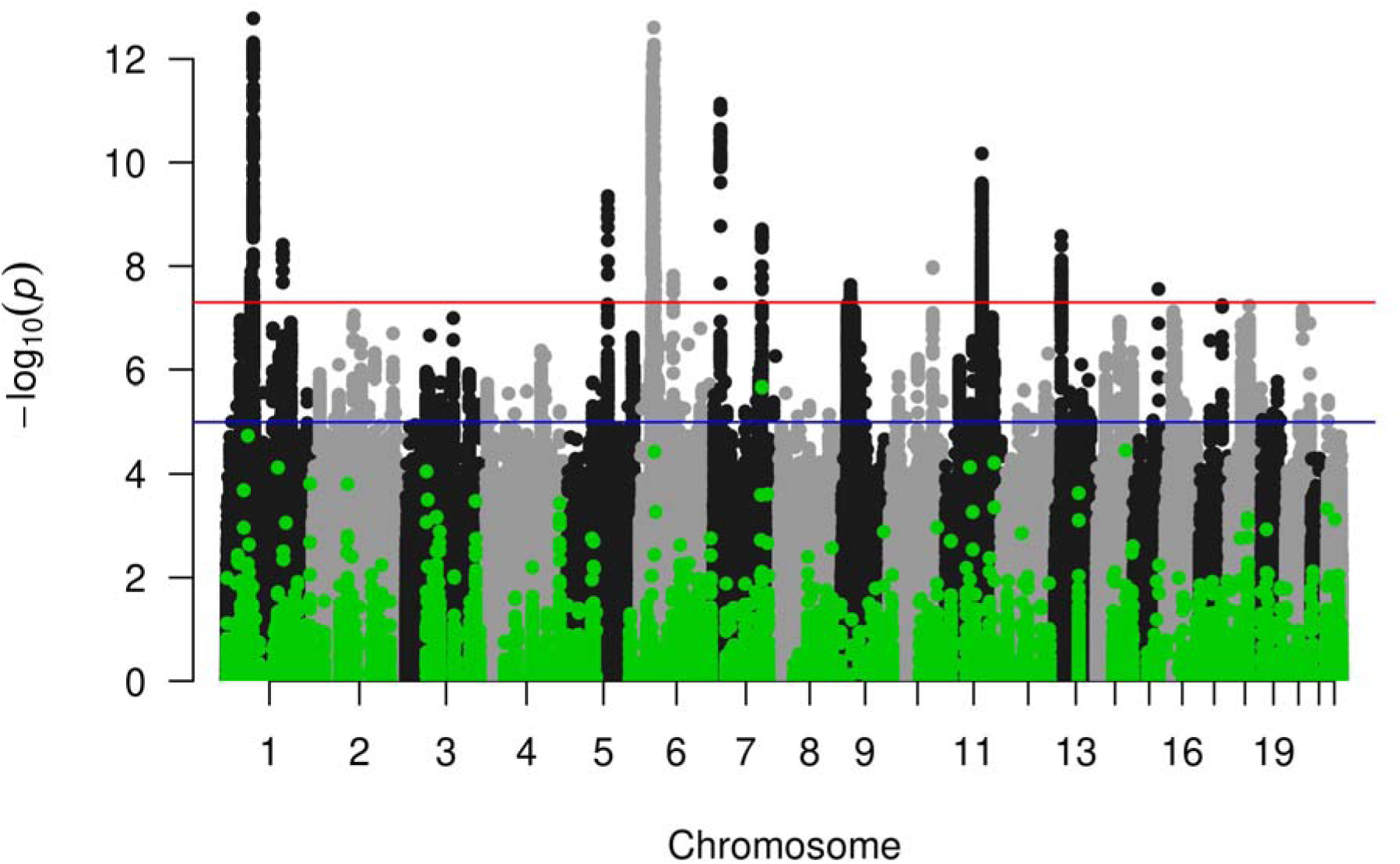
Manhattan plot for Howard (2019) broad depression GWAS results and *SLC6A3* PFC ePGS SNPs. Gray and black dots represent -log10(p) from the broad depression GWAS. Green dots represent -log10(p) from GTEx for the SNPs included in *SLC6A3* PFC ePGS. It demonstrates that all SNPs from the ePGS are not statistically significant at the genome wide level.

The results of analyses using the ePGS method have consistently revealed significant and gene x environment interactions. What is essential to appreciate is the high degree of replication of these findings across highly diverse populations, including those of different ancestry. There are now a number of published studies that demonstrate the capacity of the ePGS to identify gene-environment interactions. Importantly, these analyses use a variety of measures of environmental quality and phenotypic outcomes. For example, De Lima et al (2022) described that PFC ePGS based on the leptin receptor gene moderated the effect of postnatal adversity on child eating behaviour^41^. This was an example of a hypothesis-driven analysis based on prior knowledge of leptin receptor activity in appetite regulation. Dalmaz et al (2021) showed that a network of genes co-expressed with the synaptic protein VAMP1 gene in the PFC moderates the influence of the early environment on cognitive function in children^42^. Miguel et al (2019) found a significant association between history of exposure to perinatal hypoxic ischemic conditions and children’s cognitive flexibility, but this was moderated by the PFC *SLC6A3* ePGS ^43^.

In a study that used a WGCNA approach to define the ePGS, Arcego et al^44^ provided evidence for a hippocampal glucocorticoid-sensitive gene network as a moderated of the effect of early life adversity on later mental health in two distinct populations. The ePGS was based on a gene network derived from RNAseq with hippocampus in non-human primates using WGCNA to identify the glucocorticoid-sensitive module. Interestingly, the authors also used parallel independent component analysis to identify brain regions significantly associated with the glucocorticoid-sensitive gene network. In sum, an increasing evidence suggests that the ePGS is an appropriate method to identify GxE interaction effects.

### ePGS has high trans-ancestry portability of genetic data

Allele frequency varies across ancestries^45^ and the lack of proper diverse populations representation in current genetic association studies hampers the translation of findings into clinical applications^46^. Efforts are being made to identify genetic variations common and unique to different populations, such as the 1000 Genomes Project that identified novel SNPs^47^ and the HapMap consortium^48^. Nevertheless the level of precision currently available for European ancestry is still not uniformly available for other ancestries^49^. In PRS, the SNP list is derived from the GWAS and the same variants are included in the calculation of the polygenic score in diverse populations, which challenges PRS trans-ancestry portability^46,50,51^. The calculation of a PRS relies on SNPs, a level of analysis at which ancestral differences are greatest. In contrast, as the ePGS calculation emerges from a gene list, the SNPs included in the same ePGS may differ across ancestries but will still represent the same gene list and the same co-expression network.

The use of genetic scores that perform functional annotation or that consider genes as the first level of information, instead of SNPs, may have advantages for trans-ancestry application of genetic data^52,53^, as is the case of the ePGS method. Indeed, we see high trans-ancestry portability and replicability of findings using ePGS^9,15–17,42,43,54^. To illustrate the differences between the traditional PRS and the ePGS in terms of score composition and trans-ancestry portability, we calculated PRSs of comparable size to ePGS (*SLC6A3* or *DRD2*) in the 1000 Genomes Project dataset. The scores were calculated separately for each ancestry to account for ancestry-specific allele frequencies and linkage disequilibrium. Ancestries include African, American, East Asian, European and South Asian (**Supplemental Methods**). The same number of SNPs present in each ePGS for each ancestry was selected from the most significant variants described in the reference GWAS (broad depression^39^), and subjected to linkage disequilibrium clumping (r^2^<0.2) for calculation of PRS separately in each ancestry. Next, the SNPs derived from the calculated PRSs for each ancestry were assigned to genes and compared with ePGSs gene list. **Figure 6** shows the gene overlap between the five different ancestries for each ePGS and their respective comparable size PRS. The ePGS has a higher percentage of gene overlap between different ancestries in comparison to PRS scores in the examples seen here. These results could explain the performance of the ePGS in terms of replication seen in studies across ancestries using the ePGS method^9,15–17,42,43^ since ePGS preserves more information (number of genes) across ancestries in comparison to PRS. We also compared the score distribution density across ancestries (**Supplemental Figure 4**). Overall, the ePGS has a greater density overlap between ancestries than the PRS.

**Figure 6.**
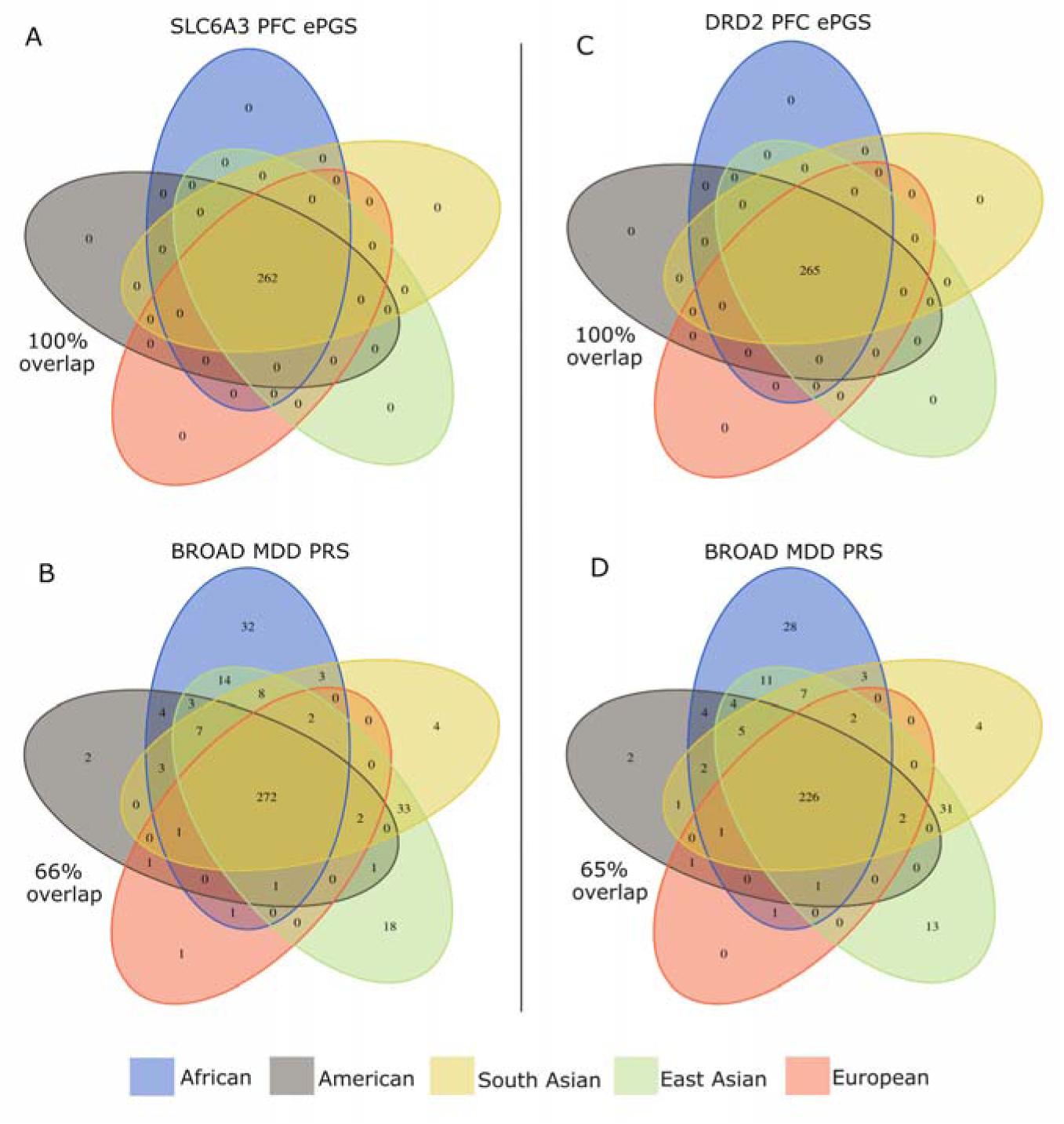
Venn diagrams of gene overlap for ePGSes and PRSes calculated based on the ePGS and PRS in the 1000 Genomes Project dataset. Gene overlap between the five different ancestries for *SLC6A3* and *DRD2* ePGS and their respective comparable size PRS. It demonstrates that the ePGS have more common genes between different ancestries in comparison to PRS scores.

### Future steps and perspectives in ePGS research

The ePGS calculation is initiated by the definition of a biologically relevant gene network, and this can be done in multiple ways. The examples provided here utilized co-expression data from mice anchored in specific genes for the identification of co-expression networks (*SLC6A3* or *DRD2*). However, other types of data and levels of information can also be used to inform the calculation of ePGS, such as protein-protein interactions, DNA methylation data, or differently expressed gene lists^17^. A promising venue currently being used in our lab consist of utilizing weighted gene correlation network analyses (WGCNA)^18^ in RNAseq data to identify co-expression gene networks significantly associated with an exposure or condition in controlled animal model experiments or in postmortem human tissue, in a data driven manner, thus completely abandoning the hypothesis-driven approach. This perspective is well aligned with the complex system in biology paradigm, and it is an anticipated improvement of the method. Arcego et al (2023) is a demonstration of this improvement as the authors used WGCNA to identify a hippocampal network of genes responsive to glucocorticoid treatment in macaques and then calculated an ePGS in humans based on this identified gene network^44^.

After the selection of the gene network, the list of genes can be filtered by diverse parameters. Adding filters allow the integration of additional information such as the developmental period, by filtering the gene selection for genes upregulated during a certain stage using Brainspan^9,34,55^. Chromosome conformation information can also be added^56^, by using data from high-throughput sequencing (Hi-C) and assigning noncoding SNPs to their cognate genes based on long-range interactions using H-MAGMA^57^ input files that describe gene–SNP pairs based on brain Hi-C data^58^. FIMO^59^ can also be used to include variants affecting transcription factor binding motifs from the genes of the network. Finally, candidate regulatory variants can be added by mapping available SNPs on promoter regions (up to 4kb upstream of the transcription start site) of the genes that compose the network. Lastly, the weight attributed to each SNP in the ePGS calculation can be derived from different GWASs. In the current examples, a GWAS for gene expression (GTEx^19^) was used, thus reflecting individual variations in gene expression of the network in the specific brain region. All these parameters can be accommodated to contemplate different research questions. Finally, adaptation of the ePGS technique for the use of single-cell and spatial transcriptomics will add still increased resolution and specificity to the polygenic scores.

## Discussion

Aligned with the idea of incorporating functional genomics information to PRS technology, we have developed the expression based polygenic score (ePGS). While both PRS and ePGS summarize the small effects of multiple SNPs using the genotype information, the use of tissue specific gene expression data in the ePGS technique transforms the polygenic score into a functional genomic tissue-specific measure. The ePGS also reflects the combined biological function of gene networks.

Here we demonstrated the consequences of rethinking SNP selection and incorporating other levels of information to polygenic scores, such as gene expression and tissue specific data. We compare ePGS and PRS features and score content. The ePGS reflects cohesive gene networks, demonstrating a high level of co-expression between the genes. This could be explained by ePGS considering only exon DNA sequences and being built from gene co-expression information. It is important to highlight that since genes do not work in isolation, but rather in networks^5^, the use of a gene network perspective has the potential to better reflect biological functions associated with these genes. We demonstrated that the ePGS and PRS reflect different biological processes, when comparing unique elements that are related to a common gene ontology term. The ePGS unique elements, in the examples demonstrated here, appear to be richer and more connected, suggesting that variations in the ePGS score may represent variation on a specific biological process. We also demonstrated that ePGS based gene networks represent tissue specific co-expression networks in humans. The possibility of reflecting functional genomics information in a tissue specific manner is one of the strengths of the ePGS, demonstrated here by the uniqueness of the *SLC6A3* PFC gene network in comparison to the *SLC6A3* Striatum gene network. As a consequence of these above-mentioned features, the ePGS is suited to test gene by environment effects, evidenced by previous published studies^9,16,42–44^.

The content of ePGS on different ancestries seem consistent when comparing the ePGS and PRS score gene overlap. This is expected since the use of genome functional annotation has the power to improve prediction of complex traits within and between ancestries^60^ and the incorporation of functional markers, such as gene expression, improves trans-ancestry portability of genomic data^61^. The ePGS uses genome functional annotation in two steps of its calculation; in the co-expression basis and by weighing the SNPs using GTEx genotype-gene expression association.

An advantage of using a gene network approach like the ePGS is the possibility of integrating other data modalities also represented by networks or with high dimensionality. For example, the integration of genetic and neuroimage information by parallel independent component analysis, which estimates the maximum independent components within each data modality separately while also maximizing the association between modalities using an entropy term based on information theory ^62^. Studies using pICA and the ePGS have found interesting results linking both data modalities and informing on the neuroanatomical basis of the effects of variations in the gene network expression^9,42,43,63^.

In conclusion, the ePGS method is purely based on biological, co-expression data and no information on association with outcomes of interest (e.g. GWAS for diseases) is used. The differences between conventional PRSs and ePGSs presented here, may explain the successful ePGS performance in gene by environment interaction models and across ancestries, suggesting that the ePGS is an interesting method to capture individual biological variation in response to environmental changes^7,17^, and may profoundly influence the way we study human disease biology.

## Acknowledgments

We thank the 1000 Genomes Project for the data availability and Sachin Patel for genetic score calculations. This research was supported by the Canadian Institutes of Health Research (CIHR, PJT-166066 and PJT-173237, PPS), Natural Sciences and Engineering Research Council of Canada (NSERC, RGPIN-2018-05063, PPS) and Fonds de recherche du Québec – Santé (FRQS to BB and PPS), awarded for the project: Le rôle de l’expression du réseau de gènes du transporteur de la dopamine sur le cerveau dans la modulation des réponses aux facteurs environnementaux au début de la vie. Relevant research was supported by the Hope for Depression Research Foundation to MJM.

## Author contributions

BB and PPS designed the study, BB, EJMF, DMA and IP conducted data analysis, BB, EJMF, DMA and IP generated the figures and BB and PPS wrote the manuscript with editing of MJM and input from all authors. PPS and MJM supervised the research. All authors read and approved the final manuscript.

## Competing interest declaration

The authors declare no competing interests.

## Notes

https://github.com/SilveiraLab/Expression-based-polygenic-scores.-Supplemental-Table-1

### Competing Interest Statement

The authors have declared no competing interest.

