## Supplemental material for "Expression based polygenic scores - A gene network perspective to capture individual differences in biological processes"

### Methods

**Supplemental Figure 1.** Network visualization comparison between ePGS for DRD2 and PRS gene networks

**Supplemental Figure 2.** Schematic overview of RNA sequencing process from tissue collection to data analysis

**Supplemental Figure 3.** SLC6A3 Striatum gene network

**Supplemental Figure 4.** ePGS and PRS distributions across 5 different ancestries from the 1000 Genomes Project

**Supplemental Table 1** – Please access to download the Supplemental Table 1 https://github.com/SilveiraLab/Expression-based-polygenic-scores.-Supplemental-Table-1 for a complete description and list of all scores used in the manuscript

### Methods

#### *1000 Genomes Project genotyping*

1000 Genomes Project[1] includes the data for 2504 unrelated participants with ancestry from 26 populations in Africa, Asia, Europe, South Asia and the Americas. All individuals were sequenced using both whole-genome sequencing (mean depth = 7.4×) and targeted exome sequencing (mean depth = 65.7×), and also genotyped using high-density SNP microarrays. A detailed description of genotyping and processing pipeline can be found in [The 1000 Genomes Project Consortium. A global reference for human genetic variation. Nature 526, 68–74 (2015). https://doi.org/10.1038/nature15393].

#### *PRS calculation – 1000 Genomes Project*

We generated polygenic scores using our accelerated pipeline PRSoS (https://github.com/MeaneyLab/PRSoS) [2], for each individual in each ancestry of the 1000 Genomes Project. The PRS were generated using broad depression GWAS[3] considering the same number of top SNPs as in the corresponding ePGS PFC SLC6A3 (12147 SNPs) or ePGS PFC DRD2 (10481 SNPs). We used the PRS for broad depression since dysfunction of the dopamine system is associated with depression symptoms[4]. First, we selected top SNPs from broad GWAS (12147 or 10481). After selection, SNPs were subjected to linkage disequilibrium clumping ($r^2$>0.2 within 500 kb radius from an index variant) separately for each ancestry. Lastly, the PRS was calculated in each ancestry as a weighted sum of the SNPs remained after the clumping procedure. The comparisons then were made between the ePGS and the PRS for broad depression of comparable size to corresponding ePGS: 1) ePGS PFC SLC6A3 was compared to broad depression PRS that was calculated based on top 12147 SNPs; 2) ePGS PFC DRD2 was compared to broad depression PRS calculated based on top 10481 SNPs. The number of SNPs to select in GWAS was chosen based on the number of SNPs included in ePGS scores (SLC6A3 and DRD2) calculated for all ancestries (**Supplemental Table 1**).

***ePGS and PRS networks properties***

The topological network structures for the genes that composed the SLC6A3 PFC ePGS, DRD2 PFC ePGS, PRS MDD comparable size with SLC6A3 PFC ePGS and PRS MDD comparable size with DRD2 PFC ePGS were visualized using Cytoscape® software[5]. To obtain gene by gene co-expression quantification we utilized GeneMANIA app (http://genemania.org)[6], selecting only connections based on co-expression. Since the GeneNetwork database provides RNA-seq information and only correlation between gene expression values of one gene against all other genes, we use the GeneMANIA database (Application version: 3.6.0), which includes curated information on genes coexpression from published papers. The information from GeneMANIA allows for the identification of all gene's interactions, while GeneNetwork does not provide such information. Cytoscape uses the information from GeneMANIA to plot the networks, giving a visual representation of the quantification of connections between the genes based in coexpression from GeneMANIA. It is important to notice that studies sourced by GeneMANIA are not tissue specific. Also, the sources used by GeneNetwork and GeneMANIA might not be identical, leading to dissimilarities in gene coexpression. Then, using the Centiscape app, inside Cytoscape, we calculated the total degree centrality measure for each gene, which reflects the number of connections a gene has with other genes. One-way ANOVA was used to verify if the difference in mean total connectivity between the networks (based on ePGS and PRS) is significant (**Figure 2C** and **Supplemental Figure 1C**). Centiscape app was also used to calculate another centrality measure, betweenness (number of times a node lies on the shortest path between other nodes). Both centrality measures were incorporated in the network visualization, the degree is represented by node size, in which bigger nodes indicate higher degree in the network visualization and betweenness is represented by node color, in which darker colors indicate higher betweenness in the networks. A control gene network was constructed using the same steps described above to visually compare the network properties with the ones described in this study in **Figure 2A** and **Supplemental Figure 1A**. The control gene network was based on a list of genes co-expressed with SLC6A3 gene in the striatum (number of genes = 346).

#### *Biological functions associated with ePGS and PRS networks*

To evaluate if the ePGS and PRS share common biological processes, we performed functional enrichment analysis for the genes that compose each network, considering a false discovery rate (FDR) adjusted p-value <0.05 to select significant gene ontology (GO) terms. The genes that compose each network were uploaded into MetaCore® software from Clarivate Analytics (https://portal.genego.com). The function "compare experiments" in MetaCore® was utilized to obtain common and unique enrichment terms between the two gene networks. We selected a significant common GO term present in both SLC6A3 and DRD2 comparisons, the term neuron differentiation. Then we built direct interaction networks inside MetaCore® using the unique elements of each network for the GO enrichment term selected to visually compare the different contribution of ePGS and PRS to the term.

#### *Assessment of gene expression patterns across human development for the ePGS and PRS associated genes*

We tested if the genes from ePGS and PRS have a notable pattern of gene expression in humans, especially for the ePGS since the co-expression data was generated from animal models. For that, we used human post-mortem gene expression samples from BrainSpan (http://brainspan.org) [7] and selected gene expression data from the genes comprising our ePGS or PRS networks. We then analyzed the correlation between expression levels for these genes in the prefrontal cortex (including the ventrolateral prefrontal cortex, orbital frontal cortex, medial prefrontal cortex, dorsolateral prefrontal cortex) from BrainSpan (from embryonic to adulthood, N= 42). Visualization of co-expression correlation matrix for each gene list was computed in R software (https://www.r-project.org) using the "cor" function and plotting it as correlation matrix. Next, we computed the percentage of pairs of genes with absolute value of expression correlation higher than 0.5 (considered a high correlation for gene expression) and the correlations significant at p-value <0.05, to numeric see the differences of co-expression across the different gene lists.

**Comparison between ancestries**

To compare the traditional PRS and the ePGS in terms of scores' composition and trans-ancestry portability we calculated the scores split by ancestry in the 1000 Genomes Project dataset (see Methods sections "Expression-based polygenic scores (ePGS or ePGS) calculation" and "PRS calculation – 1000 Genomes Project" and Results section) and compared the PRS and ePGS for each ancestry group in two ways. 1) SNP included in PRS and ePGS were converted to genes using the biomaRt R package[8,9], and we compared the sets of genes between PRS and ePGS using the VennDiagram R package[10]. The percentage of overlap between the PRS and ePGS distributions was calculated as the number of genes in the overlap divided by the total number of unique genes included in PRS and ePGS. 2) We also compared the score distributions across ancestries. For that, we approximated distributions of the PRS and ePGS with kernel density and calculated the overlap as the proportion of the area that is overlapped to the total area.

**Supplemental Figure 1. Network visualization comparison between ePGS and PRS for DRD2 networks**

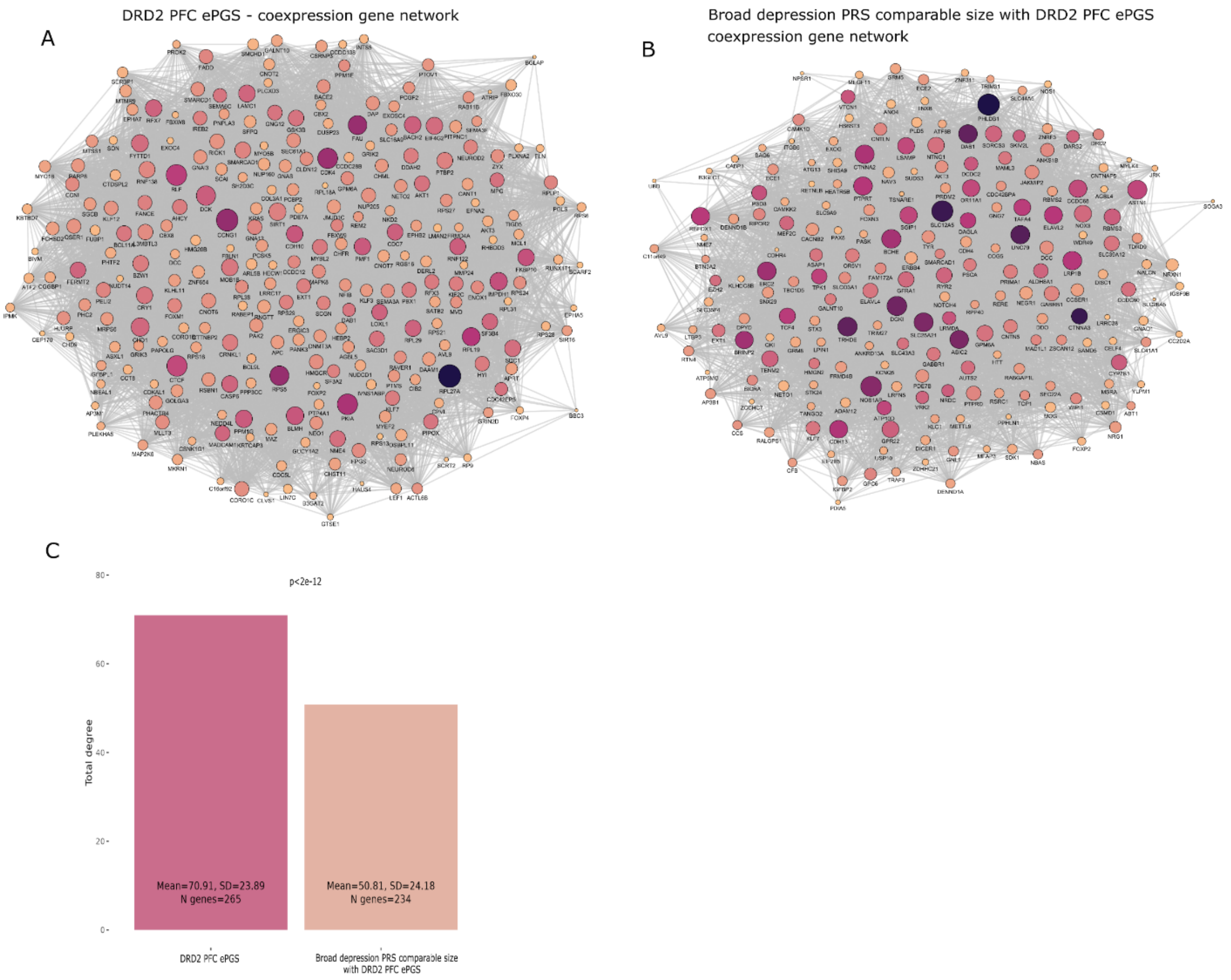


**Supplemental Figure 1 – DRD2 PFC ePGS coexpression gene network and the respective comparable size broad depression PRS. a)** DRD2 PFC ePGS gene network; **b)** broad depression PRS gene network comparable size with DRD2 PFC ePGS; **c)** one-way ANOVA results of total connectivity comparison (total degree values) for DRD2 ePGS and PRS comparable size. Gene co-expression interactions were obtained from GeneMANIA ([http://genemania.org](http://genemania.org)) and used to generate the networks with Cytoscape® application, which specifies amount of interactions between pairs of genes based on their co-expression, represented by the number of edges (gray lines) in the networks. The Centiscape plug-in in Cytoscape® was used to calculate the centrality of the genes in each network, defining the degree (number of connections with other nodes, represented by node size, in which bigger nodes indicates more connections with other nodes) and betweenness (number of times a node lies on the shortest path between other nodes, represented by node's color in which darker colors indicate higher betweenness in the networks) for the components of the networks.

### Supplemental Figure 2 – Overview of RNA sequencing process and data analysis

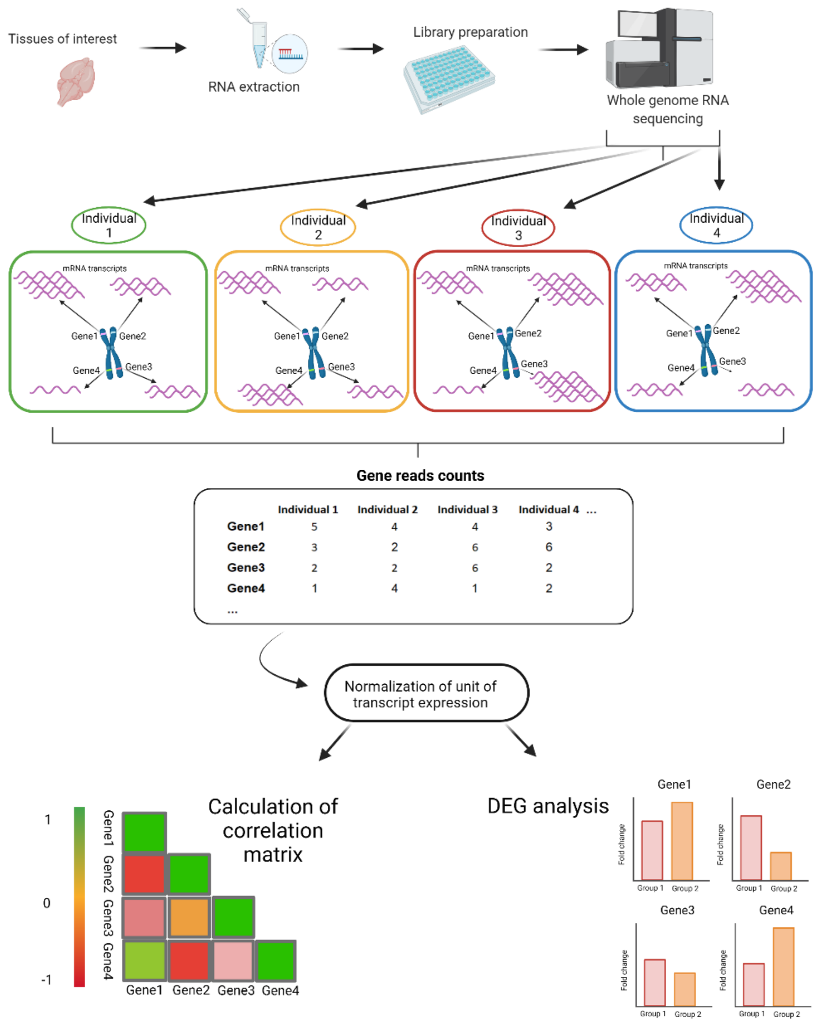


**Supplemental Figure 2. Schematic overview of RNA sequencing process: from tissue collection to data analysis.** Upper panel - a tissue of interest in chosen and collected, RNA extraction, library preparation and then whole genome RNA sequencing are performed. The process of whole genome RNA sequencing consists of identifying the amount of mRNA in each mapped gene, in each chromosome, in the whole genome. As the result, the gene read counts are obtained for each sample and for all genes mapped in the whole genome sequencing. These units of transcript expression are then normalized, by transforming in reads per kilo base per million mapped reads (RPKM) to account for variations in genes' length. RPKM can be analyzed in several ways. Here are represented two possible analyses. On the right is the gene differential expression analysis which compares expression of a single gene between the groups to identify the genes that are significantly different between the groups in terms of expression (differentially expressed genes, DEG). On the left is a correlation matrix between all genes that were sequenced, which will inform which genes are co-expressed together, meaning that the expression of these genes is varying together, both in a positive or negative way. Co-expression, in the present study, is used as an indicative of how much these genes are working together, thus possibly in the same biological process.

**Supplemental Figure 3. SLC6A3 Striatum gene network**

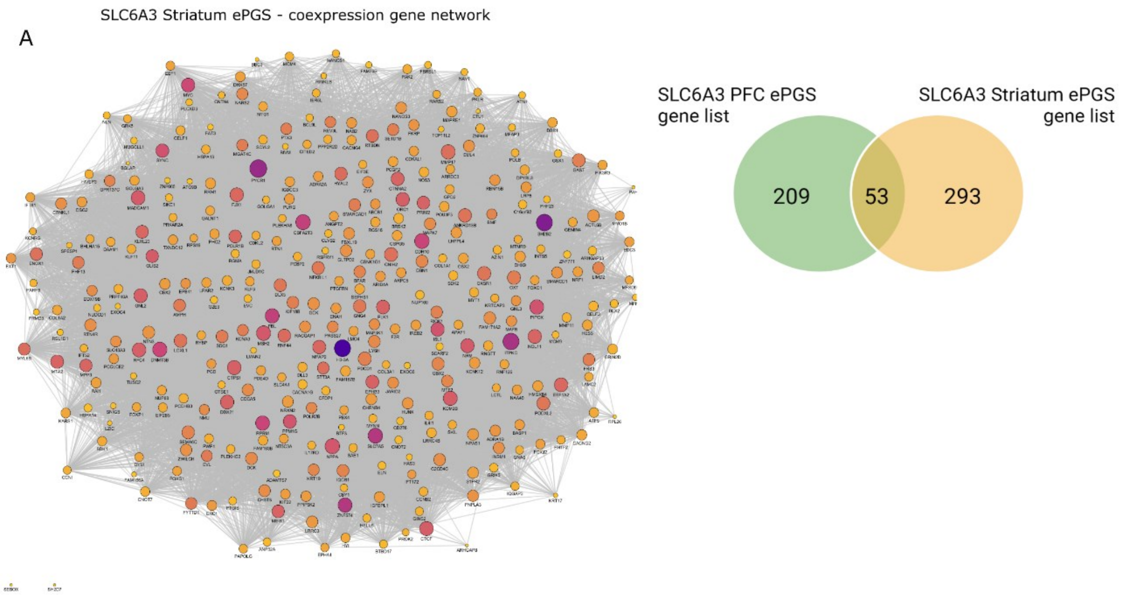


**Supplemental Figure 3 –Visualization for the SLC6A3 Striatum ePGS gene network. a)** SLC6A3 Striatum ePGS coexpresion gene network. Gene co-expression interactions were obtained from GeneMANIA (http://genemania.org) and used to generate the networks with Cytoscape® application, which specifies the amount of interactions between pairs of genes based on their co-expression, represented by the number of edges (gray lines) in the networks. The CentiScaPe plug-in[11] in Cytoscape® was used to calculate the centrality of the genes in each network, defining the degree (number of connections with other nodes, represented by node size, in which bigger nodes indicates more connections with other nodes) and betweenness (number of times a node lies on the shortest path between other nodes, represented by node's color, in which darker colors indicate higher betweenness in the networks) for the components of the networks; **b)** Gene overlap between SLC6A3 PFC ePGS gene list and SLC6A3 Striatum ePGS gene list.

### Supplemental Figure 4. ePGS and PRS score distribution across 5 different ancestries

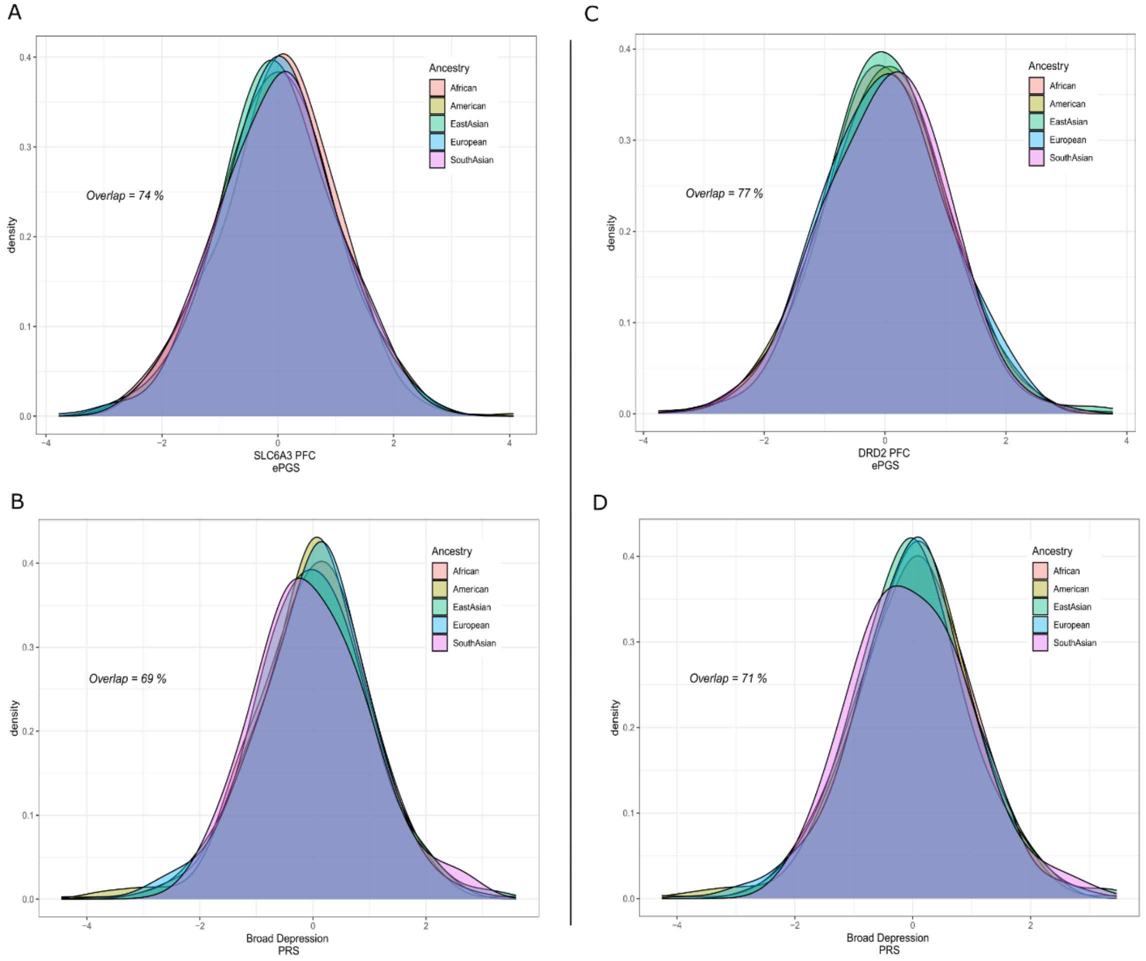


**Supplemental Figure 4 –Visualization of scores distribution overlap across 5 different ancestries.** Scores for each ancestry were z-transformed and distributions were plotted and then compared. **a)** ePGS SLC6A3 PFC score; **b)** Broad depression PRS of comparable size with ePGS SLC6A3 PFC score; **c)** ePGS DRD2 PFC score; and **d)** broad depression PRS of comparable size with ePGS DRD2 PFC score.
